# A specific non-bisphosphonate inhibitor of the bifunctional farnesyl/geranylgeranyl diphosphate synthase in malaria parasites

**DOI:** 10.1101/134338

**Authors:** Jolyn E. Gisselberg, Zachary Herrera, Lindsey Orchard, Manuel Llinás, Ellen Yeh

## Abstract

Isoprenoid biosynthesis is essential for *Plasmodium falciparum* (malaria) parasites and contains multiple validated antimalarial drug targets, including a bifunctional farnesyl and geranylgeranyl diphosphate synthase (FPPS/GGPPS). We identified MMV019313 as an inhibitor of *Pf*FPPS/GGPPS. Though *Pf*FPPS/GGPPS is also inhibited by a class of bisphosphonate drugs, MMV019313 has significant advantages for antimalarial drug development. MMV019313 has superior physicochemical properties compared to charged bisphosphonates that have poor bioavailability and strong bone affinity. We also show that it is highly selective for *Pf*FPPS/GGPPS and showed no activity against human FPPS or GGPPS. Inhibition of *Pf*FPPS/GGPPS by MMV019313, but not bisphosphonates, was disrupted in an S228T variant, demonstrating that MMV019313 and bisphosphonates have distinct modes-of-inhibition against *Pf*FPPS/GGPPS. Altogether MMV019313 is the first specific, non-bisphosphonate inhibitor of *Pf*FPPS/GGPPS. Our findings uncover a new small molecule binding site in this important antimalarial drug target and provide a promising starting point for development of *Plasmodium*-specific FPPS/GGPPS inhibitors.

## Introduction

There is an urgent need for antimalarials with novel mechanisms-of-action to circumvent resistance to frontline drugs. The biosynthesis of cellular isoprenoids is an essential process in *Plasmodium* parasites that cause malaria. A number of antimalarial compounds target enzymes in isoprenoid biosynthetic pathways leading to parasite growth inhibition. First, *Plasmodium* parasites depend on the 7-enzyme prokaryotic 2-*C*-methyl-D-erythritol 4-phosphate (MEP) pathway in its plastid organelle, the apicoplast, to produce isopentenyl pyrophosphate (IPP) and its isomer dimethyallyl pyrophosphate (DMAPP) (Jomaa et al., 1999). IPP and DMAPP are the C5 building blocks for all isoprenoids. The antibiotic fosmidomycin inhibits the MEP enzyme, Dxr/IspC, in both bacteria and *Plasmodium* parasites (Jomaa et al., 1999).

Second, at least three isoprenoid synthases (PF3D7_1128400.1, PF3D7_0202700, PF3D7_0826400) catalyze the condensation of IPP and DMAPP into longer prenyl chains (Artz et al., 2011; Jordão et al., 2013; Tonhosolo et al., 2005). In particular, farnesyl pyrophosphate synthase (FPPS) and geranylgeranyl pyrophosphate synthase (GGPPS) are key branch point enzymes that synthesize C15 and C20 prenyl chains, respectively, for multiple downstream enzymes. In *Plasmodium* parasites, these reactions are catalyzed by a single bifunctional enzyme, the farnesyl/geranylgeranyl diphosphate synthase (Artz et al., 2011; Jordão et al., 2013). Nitrogen-containing bisphosphonates, blockbuster drugs which inhibit human FPPS, also inhibit the bifunctional *Plasmodium* FPPS/GGPPS (Ghosh et al., 2004; Jordão et al., 2011; Michael B Martin et al., 2001; No et al., 2012; Singh et al., 2010).

Finally, prenyl chains are cyclized and/or conjugated to small molecule and protein scaffolds by a variety of prenyltransferases to biosynthesize final isoprenoid products required for parasite growth and replication. Tetrahydroquinolines (THQ) have been shown to potently inhibit the *Plasmodium* protein farnesyltransferase (Eastman et al., 2005; 2007; Nallan et al., 2005). Other inhibitors may interfere with isoprenoid biosynthesis indirectly by disrupting transporters that supply starting substrates or export products or blocking pathways that provide cofactors for isoprenoid biosynthetic enzymes.

Importantly fosmidomycin, bisphosphonates, and tetrahydroquinolines have all shown efficacy in mouse models of malaria infection, validating the key importance of isoprenoid biosynthesis as an antimalarial drug target (Jomaa et al., 1999; Nallan et al., 2005; No et al., 2012; Singh et al., 2010). Fosmidomycin is currently being tested in human clinical trials, while a THQ lead candidate was investigated in preclinical studies (Fernandes et al., 2015; Nallan et al., 2005). However, novel chemical scaffolds that disrupt isoprenoid biosynthetic pathways in *Plasmodium* remain desirable to overcome unfavorable drug properties of these known inhibitors. For example, bisphosphonates avidly bind bone mineral, and both fosmidomycin and THQs have short half-lives *in vivo* (Sinigaglia et al., 2007; Tsuchiya et al., 1982; Van Voorhis et al., 2007).

In 2011 the Medicine for Malaria Venture (MMV) distributed the Open-Access Malaria Box to accelerate antimalarial drug discovery (Spangenberg et al., 2013). The Malaria Box consists of 400 structurally diverse compounds, curated from >20,000 hits generated from large-scale screens, that inhibit the growth of blood-stage *Plasmodium falciparum* parasites (Gamo et al., 2010; Guiguemde et al., 2010; Rottmann et al., 2010). A major goal of sharing these compounds was to facilitate elucidation of their antimalarial mechanism-of-action and open new classes of validated chemical scaffolds and drug targets. Compounds that disrupt isoprenoid metabolism can be detected by “rescue” of their growth inhibition upon supplementation of isoprenoids in the growth media (Yeh and Derisi, 2011). Previously, we and two other groups screened the Malaria Box for compounds whose growth inhibition were rescued by addition of IPP and identified MMV008138 (Bowman et al., 2014; Wu et al., 2015). We and our collaborators demonstrated that MMV008138 inhibits IspD, an enzyme in the MEP pathway that produces IPP (Wu et al., 2015).

Using a quantitative high-throughput screen (qHTS), we report a second compound in the Malaria Box, MMV019313, that shows an IPP rescue phenotype but was not identified in screens performed by other groups (Bowman et al., 2014; DeRisi, 2014; Van Voorhis et al., 2016). We demonstrate that the target of MMV019313 is the *P. falciparum* FPPS/GGPPS, a cytosolic isoprenoid synthase that utilizes IPP and the key branch point enzyme in isoprenoid biosynthesis in parasites. MMV019313 represents the first new class of non-bisphosphonate inhibitors of *Pf*FPPS/GGPPS.

## Results

### A quantitative high-throughput screen (qHTS) for growth and IPP rescue identifies MMV019313 as an inhibitor of isoprenoid biosynthesis

Previous IPP rescue screens of the Malaria Box tested compounds at a single, high concentration >5 µM (Bowman et al., 2014; DeRisi, 2014; Van Voorhis et al., 2016). While testing compounds at a single concentration is useful for identifying phenotypes that occur at a threshold concentration (e.g. growth inhibition), growth rescue is expected to occur within a specific range of concentrations. For example, doxycycline inhibits *P. falciparum* growth with an EC_50_= 0.3 µM that increases to 3.2 µM upon addition of IPP (Yeh and Derisi, 2011). At concentrations <0.3 µM, doxycycline does not cause growth inhibition. However, at concentrations >3.2 µM, it is no longer specific for its target and causes growth inhibition through additional targets that cannot be IPP rescued. Therefore the concentration range in which IPP rescue can be observed is greater than the EC_50_ of the compound for its specific, IPP-rescuable target but less than that for any nonspecific targets.

To increase the sensitivity for detecting IPP chemical rescue, we screened the Malaria Box for growth inhibition of blood-stage *P. falciparum* in the presence and absence of IPP over 8-12 drug concentrations from 0.01-27 µM (Table S1). Of 397 compounds tested (3 compounds were not available), 383 showed growth inhibition at ≤27 µM. Initial hits showing IPP rescue of growth inhibition at one or more drug concentrations were commercially sourced and retested. Along with the previously reported compound, MMV008138, we confirmed a second compound in the Malaria Box showing an IPP rescue phenotype, MMV019313 (Figure 1A). MMV019313 inhibited *P. falciparum* growth measured in a single replication cycle with EC_50_=268 nM (250-289 nM) in the absence of IPP; in the presence of IPP (200 µM), the EC_50_ was over 13-fold more at 3.6 µM (3.2-4.0 µM) (Figure 1B). Notably, at concentrations >3.6 µM, MMV019313 inhibits a nonspecific target that can no longer be IPP rescued, which explains why it was not identified in previous screens.

**Figure 1.**
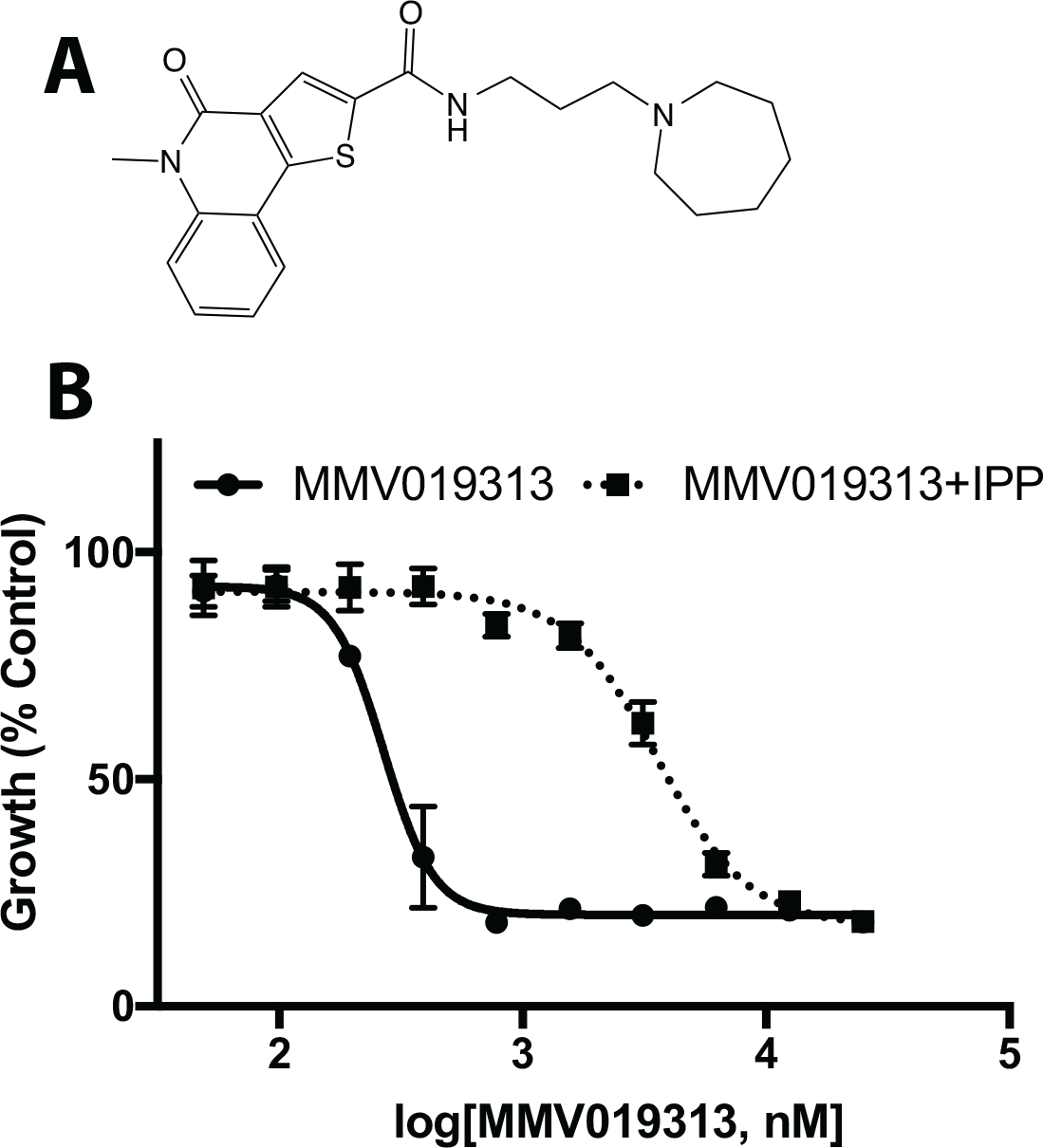
IPP rescues growth inhibition by MMV019313. **A.** The structure of MMV019313. **B.** EC_50_ curves in the absence (solid line) and presence (dotted line) of IPP. Parasitemia is normalized to that of an untreated control. Results shown for experiments performed in biological triplicate with technical duplicate. Shown plotted as mean ±SD.

An unusual feature of blood-stage *Plasmodium* is that IPP can rescue complete loss of the apicoplast, the plastid organelle which houses the MEP pathway, since production of IPP is the only essential function of the apicoplast. Compounds like doxycycline that disrupt apicoplast biogenesis cause growth inhibition rescued by IPP and result in parasites lacking an apicoplast (Yeh and Derisi, 2011). In contrast, inhibitors of isoprenoid biosynthesis, like fosmidomycin and MMV008138, cause growth inhibition rescued by IPP with an intact apicoplast (Amberg-Johnson et al., 2017). We determined whether MMV019313 disrupted the biogenesis of the apicoplast. Both the replication of the apicoplast genome and import of an apicoplast-targeted GFP were intact in MMV019313-treated and IPP-rescued parasites (Figure S1) (Yeh and Derisi, 2011). Altogether MMV019313 causes growth inhibition rescued by IPP with no defect in apicoplast biogenesis, suggesting that, like fosmidomycin and MMV008138, MMV019313 blocks isoprenoid biosynthesis (Wu et al., 2015).

### MMV019313-resistant parasites contain a mutation in the farnesyl/geranylgeranyl diphosphate synthase

Though metabolomic profiling of MMV019313 has been performed, isoprenoid intermediates were not examined so a specific disruption of these pathways could not be inferred (Allman et al., 2016; Creek et al., 2016). Instead to gain further insight into the mechanism-of-action of MMV019313, we identified mutations that confer resistance to MMV019313. Our initial attempt to select drug-resistant parasites from a bulk culture of 10^10^ parasites was not successful, though this same protocol was effective in selecting drug resistance against MMV008138 when performed in parallel (Wu et al., 2015). Therefore, we turned to chemical mutagenesis to increase the likelihood of selecting MMV019313-resistant parasites. Chemical mutagenesis followed by whole genome sequencing to identify mutations has been successfully employed to perform phenotypic screens in *T. gondii* but has not commonly been used in *Plasmodium* parasites (Desai et al., 2005; Farrell et al., 2014; Inselburg, 1984; 1985). We treated 10^8^ *P. falciparum* W2 parasites with sub-lethal doses of ethyl methanesulfonate (EMS), an alkylating agent. Because EMS is unstable, we tested a fresh batch of mutagen immediately prior to treatment in a standard parasite growth inhibition assay to determine its EC_50_. We then selected the EC_50_ (2.025 mM) as the highest EMS concentration used and tested several lower concentrations by serially diluting 3-fold. These relatively low concentrations were selected to maximize the frequency of drug-resistant mutations while minimizing lethal mutations that would result in a smaller pool of starting parasites or the presence of nonspecific passenger mutations that would confound identification of causative mutations during whole genome analysis. Following mutagenesis, parasites were continuously selected with a dose of MMV019313 equal to its EC_75_ in the presence of IPP for 32 days before removal of the drug. This concentration was chosen to maximize selection pressure for developing resistance in the IPP-rescuable target, while minimizing that for developing resistance in nonspecific targets.

Resistant parasites emerged in all mutagenized cultures one week after drug removal. In these resistant populations, MMV019313 showed EC_50_ values that ranged from 3-9-fold greater than that observed in the initial susceptible population. Growth inhibition of two MMV019313-resistant populations which showed the highest levels of resistance at the lowest EMS concentrations used (8.3 µM and 25 µM, designated 019313R1 and 019313R2 respectively) are shown in Figure 2A. Significantly the EC_50_ of MMV019313 in 019313R1 and 019313R2 was similar to that observed in susceptible strains in the presence of IPP, and growth inhibition could no longer be rescued by addition of IPP (Figure S2). These results suggest that, as expected, 019313R1 and 019313R2 populations were completely resistant to inhibition of its specific IPP-rescuable target but had not developed resistance to inhibition of additional nonspecific targets. This is the first example of using chemical mutagenesis to aid selection of drug-resistant parasites in *P. falciparum*.

**Figure 2.**
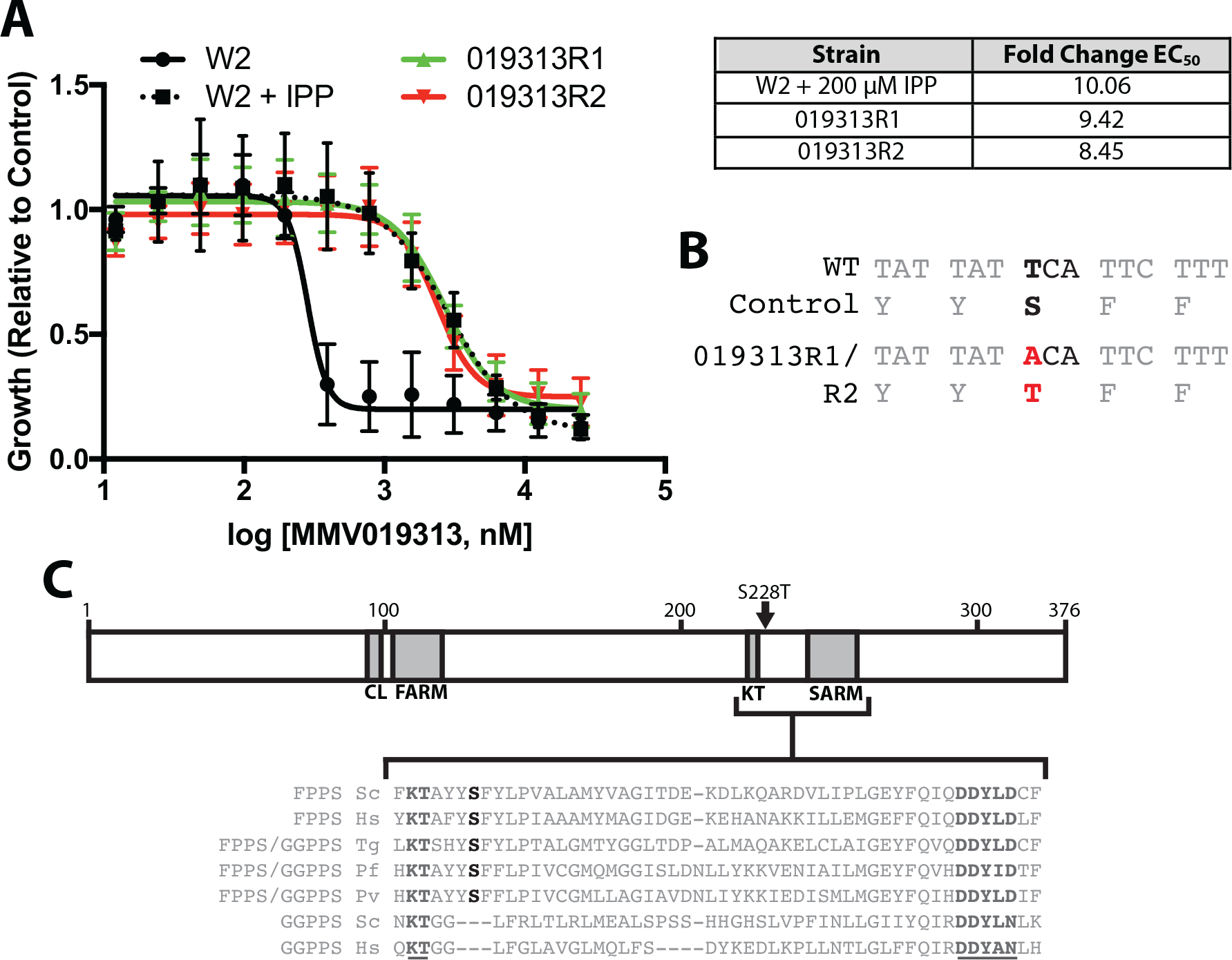
MMV019313-resistant parasites contain a mutation in the bifunctional farnesyl and geranylgeranyl diphosphate synthase. **A.** EC_50_ curves of parental W2 parasites alone (black, solid line) and with IPP supplementation (black, dotted line) and the two resistant populations (019313R1, red line, and 019313R2, green line) without IPP added. Fold change in EC_50_ compared to W2 parasites is shown in the right. Results shown for experiments performed in biological triplicate with technical duplicate. Shown plotted as mean ±SD. **B.** Mutation determined by whole genome sequencing of resistant populations 019313R1 and 019313R2, highlighted in red **C.** Schematic of *Pf*FPPS/GGPPS protein. The S228T residue is highlighted in bold black while conserved KT and SARM residues are highlighted in bold grey and underlined.

019313R1, 019313R2, and their respective mutagenized parent strains used to initiate drug selection, WT1 and WT2, were subjected to whole-genome sequencing. Comparison of the resistant and parent genome sequences identified a single nucleotide variant (SNV) common to both resistance populations, which was present in 100% of reads in 019313R1 and 63% of reads in 019313R2 but not present in either susceptible parent genomes (Table 1). No other SNV were detected at >40% prevalence in either resistant populations relative to the corresponding parent populations (Table S2). The identified SNV was a T-to-A mutation in the gene PF3D7_1128400 characterized as a bifunctional FPPS/GGPPS and resulted in a Ser228-to-Thr change in the protein (Figure 2B) (Artz et al., 2011; Jordão et al., 2013). IPP is a known substrate of the FPPS/GGPPS, and in the primary sequence Ser228 is adjacent to conserved KT residues and an Asp-rich region required for catalysis in all homologous FPPS and GGPPS enzymes (Figure 2C). *Pf*FPPS/GGPPS was a strong candidate for further validation as the molecular target of MMV019313.

**Table 1.** Summary of mutations identified by whole genome sequencing

| Gene ID<sup>a</sup> | Description<sup>b</sup> | Base Call<sup>c</sup> |  |  | AA change | Population | Read Number<sup>d</sup> |  | %<sup>e</sup> |  |
| --- | --- | --- | --- | --- | --- | --- | --- | --- | --- | --- |
|  |  | Position | WT | EMS |  |  | WT | R | WT | R |
| PF3D7_1128400 | Geranylgeranyl pyrophosphate synthase (GGPPS) | 682 | T | A | S228T | 019313R1 | 219 | 163 | 100 | 100 |
|  |  |  |  |  |  | 019313R2 | 186 | 181 | 100 | 62.4 |

<sup>a</sup> PlasmoDB gene Identification number
<sup>b</sup> Basic gene description based on PlasmoDB functional assignments
<sup>c</sup> WT calls match 3D7 reference genome
<sup>d</sup> Read depth corresponding to WT call (IGV) and Mut call (SnEff) respectively
<sup>e</sup> Percent of reads corresponding to WT call (IGV) and Mut call (SnEff) respectively

a PlasmoDB gene Identification number

b Basic gene description based on PlasmoDB functional assignments

c WT calls match 3D7 reference genome

d Read depth corresponding to WT call (IGV) and Mut call (SnpEff) respectively

e Percent of reads corresponding to WT call (IGV) and Mut call (SnpEff) respectively

### Overexpression and an S228T variant of *Pf*FPPS/GGPPS confers resistance to MMV019313

We determined whether overexpression of wildtype *Pf*FPPS/GGPPS was sufficient to confer resistance to MMV019313. A transgene encoding FPPS/GGPPS-GFP under the control of either the ribosomal L2 protein (*RL2*; PF3D7_1132700) or calmodulin (*CaM*; PF3D7_1434200) promoter was integrated into an engineered *att*B locus in Dd2^attb^ parasites (Nkrumah et al., 2006). Expression of the CaM promoter is 10 to 50-fold greater across the life cycle than that of the RL2 promoter based on RNA-seq data, resulting in appreciably greater FPPS/GGPPS-GFP protein (Figure S3A) (Otto et al., 2010). Therefore, we compared the effect of no, moderate (RL2), and high (CaM) levels of FPPS/GGPPS-GFP overexpression on susceptibility to MMV019313. As shown in Figure 3A and Table S3, overexpression of FPPS/GGPPS-GFP results in a dose-dependent increase in the *EC*_50_ of MMV019313 with intermediate-level resistance to MMV019313 observed in the RL2-FPPS/GGPPS-GFP strain and high-level resistance observed in the CaM-FPPS/GGPPS-GFP strain. Notably growth inhibition by MMV019313 in the CaM-FPPS/GGPPS-GFP strain could no longer be rescued by IPP, indicating that complete resistance to inhibition of the specific IPP-rescuable target had been achieved (Figure S3B).

**Figure 3.**
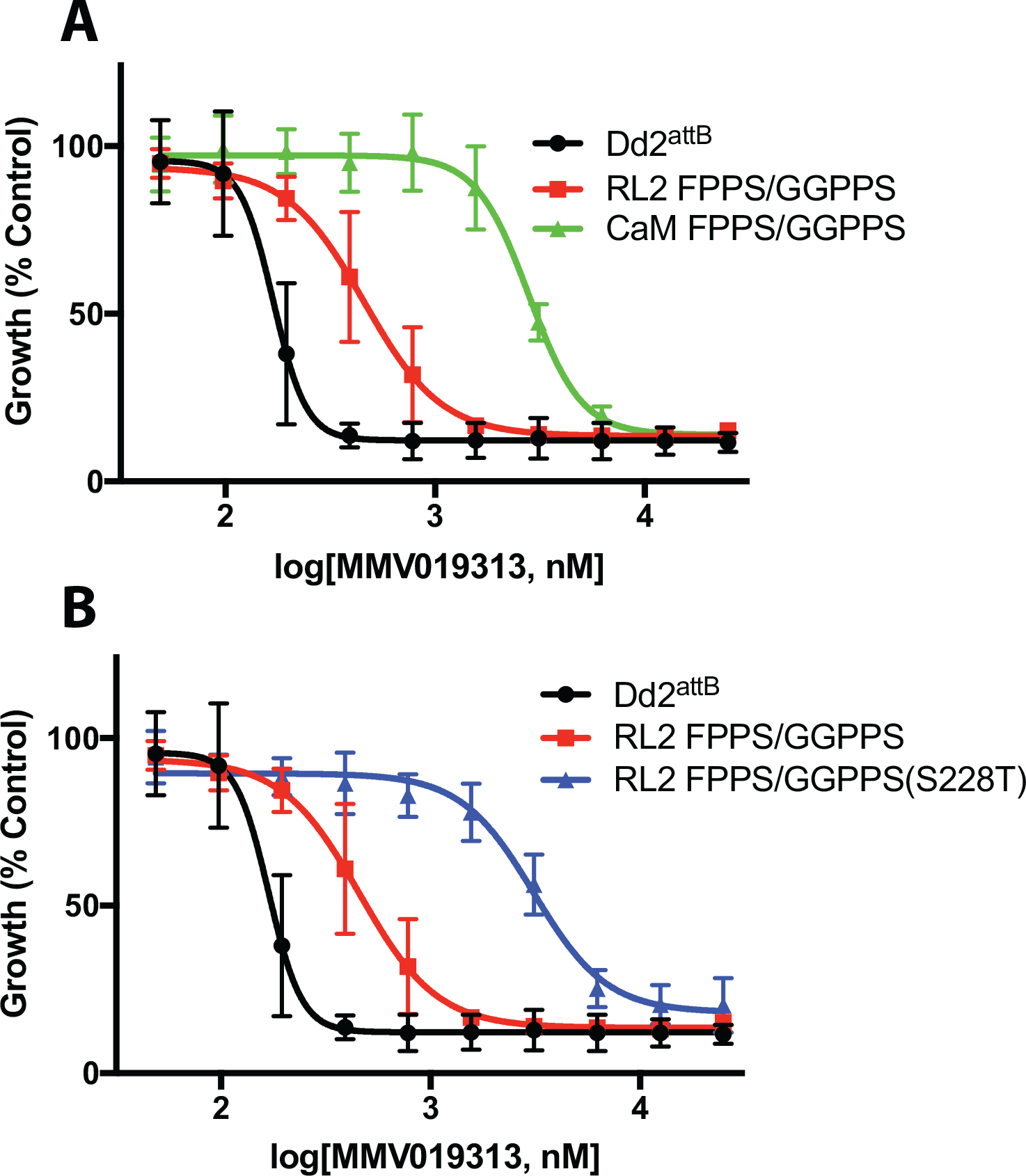
Overexpression of WT and an S228T variant of *Pf*FPPS/GGPPS confer resistance to MMV019313. **A.** EC_50_ curves of MMV019313 against the parental Dd2^attB^ parasites (black) and parasites over-expressing FPPS/GGPPS-GFP under the RL2 (red) or CaM (green) promoter. **B.** EC50 curves of MMV019313 against the parental Dd2^attB^ parasites (black) and parasites over-expressing wild type (red) or mutant (S228T, blue) FPPS/GGPPS-GFP under the RL2 promoter. Results shown for experiments performed in biological duplicate with technical duplicate. Shown plotted as mean ±SD.

We also determined whether the S228T variant of GGPPS confers MMV019313 resistance. Initially we attempted to introduce the S228T mutation into the *fpps/ggpps* gene in the *P. falciparum* genome using CRISPR-Cas9 mutagenesis but were unable to recover mutant parasites. As an alternative, we overexpressed the FPPS/GGPPS(S228T)-GFP variant using the “moderate” RL2 promoter and compared its effect on MMV019313 susceptibility with that of the wild-type RL2-FPPS/GGPPS-GFP. Overexpression of the FPPS/GGPPS(S228T)-GFP variant, even at moderate levels, caused an 18-fold increase in the *EC*_50_ which did not increase further with IPP rescue, indicating that complete resistance to inhibition of the specific, IPP-rescuable target had been achieved (Figure 3B and S3C; Table S3). Because moderate overexpression of the S228T variant caused greater MMV019313 resistance than moderate overexpression of wildtype *Pf*FPPS/GGPPS, we conclude that the S228T variant is sufficient to confer resistance independent of overexpression. Altogether these results clearly demonstrated that the mechanism-of-action of MMV019313 is dependent on *Pf*FPPS/GGPPS.

### MMV019313 specifically inhibits the enzymatic activity of *Pf*FPPS/GGPPS

To confirm that *Pf*FPPS/GGPPS is the molecular target of MMV019313, we directly measured MMV019313 inhibition in enzymatic assays. Consistent with a previous report, purified *Pf*FPPS/GGPPS catalyzed the production of both farnesyl(C15)-PP and geranylgeranyl(C20)-PP (Jordão et al., 2013). To measure the inhibitory effect of MMV019313, we determined its *IC*_50_ for FPP and GGPP production in two conditions. In the first “non-saturating” condition, substrate concentrations equaled *K*_M_ values (Figure S4), the concentration at which the rate of reaction is half-maximal. In the second “*k*_cat_” condition, substrate concentrations were saturating and the rate of reaction was maximal. Measuring inhibitor effects in both “non-saturating” and “*k*_cat_” conditions increases the sensitivity for detecting different types of inhibitors (competitive, non-competitive, uncompetitive). We found the *IC*_50_ value was 330 nM for FPP production under “non-saturating” conditions, comparable to its *EC*_50_ value in cellular growth inhibition assays (Figure 1B and S5). MMV019313 was identified by its IPP rescue phenotype in cell growth assays wherein addition of exogenous substrate increases enzymatic rates, indicating that physiological conditions are likely sub-saturating. Therefore its *IC*_50_ value measured in sub-saturating condition is more relevant for comparison to its *EC*_50_ value for growth inhibition. Under “*k*_cat_” conditions, *IC*_50_ values were 2.0 µM for FPP production and 9.8 µM for GGPP production (Figure 4A-B).

**Figure 4.**
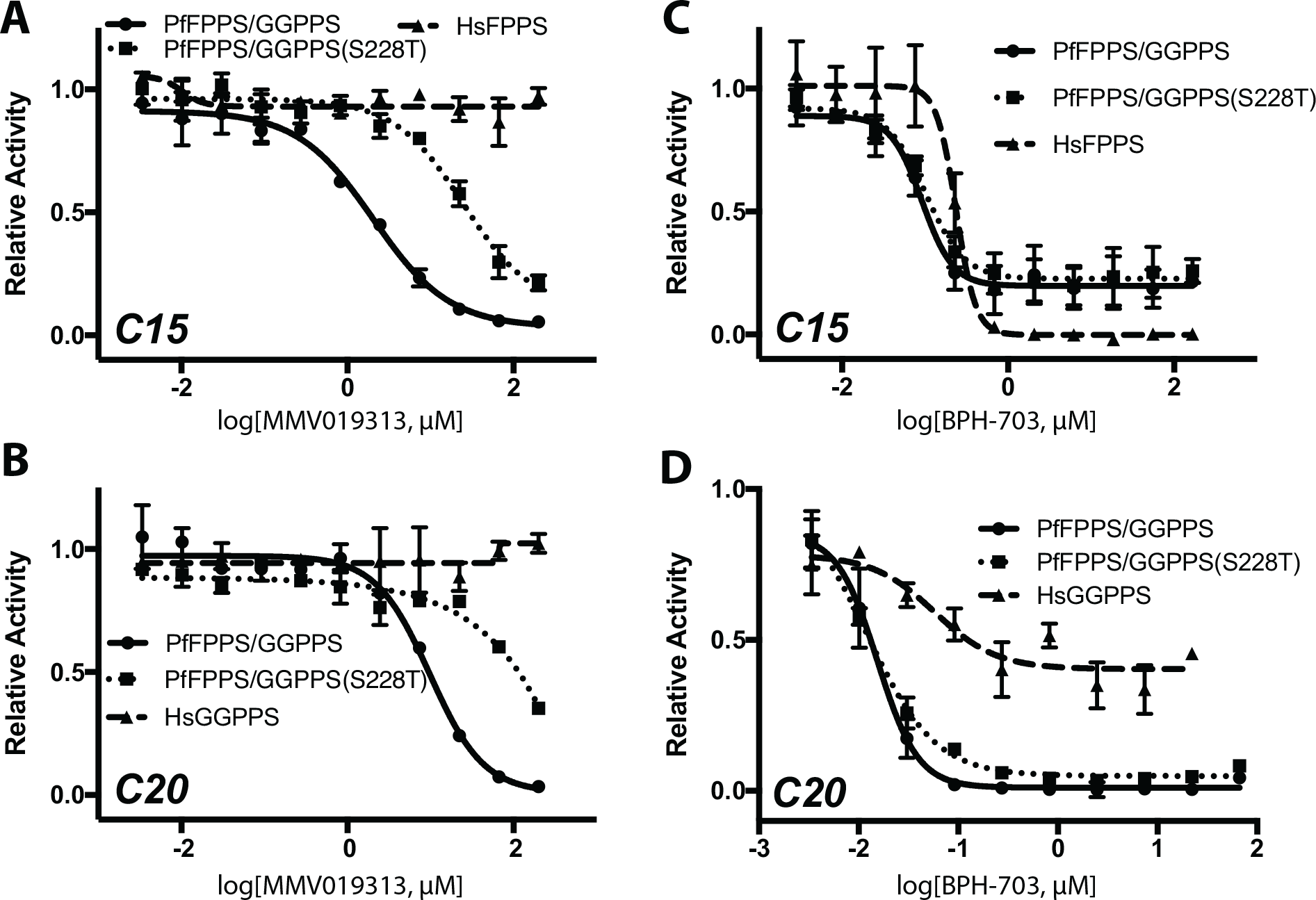
MV019313 has a specific and distinct mode-of-inhibition against purified *Pf*FPPS/GGPPS. Dose-dependent inhibition of wild-type *Pf*FPPS/GGPPS (solid), *Pf*FPPS/GGPPS S228T variant (dotted), and either human FPPS or GGPPS (dashed) of **A**. FPP (C15) production by MMV019313, **B**. GGPP (C20) production by MMV010313, **C**. FPP (C15) production by BPH-703, and **D**. GGPP (C20) production by BPH-703. Partial inhibition of *Hs*GGPPS by BPH-703 was observed under *k*_cat_ conditions, while complete inhibition was observed under non-saturating conditions (Figure S5). Shown plotted as mean ±SD.

Since inhibition was detectable in both conditions, assays of mutant and human enzymes were performed at saturating conditions which gives higher signal-to-noise. Significantly MMV019313 at concentrations up to 200 µM did not inhibit human FPPS or GGPPS activity, indicating that it was at least 100-fold selective for *Pf*FPPS/GGPPS over human homologs (Figure 4A-B; Table S3). In contrast, the most selective bisphosphonate identified by a previous study, BPH-703, showed only a 2.6-fold selectivity for *Pf*FPPS/GGPPS versus human FPPS and 3.6-fold selectivity versus human GGPPS in enzymatic assays (Figure 4C-D; Table S3) (No et al., 2012).

### MMV019313 binds a novel site on *Pf*FPPS/GGPPS, distinct from that of bisphosphonates

The lack of inhibition of human FPPS and GGPPS by MMV019313 was intriguing since *Pf*FPPS/GGPPS is structurally related to human FPPS and GGPPS and all are inhibited by bisphosphonates (No et al., 2012). Co-crystal structures have shown that bisphosphonates, zoledronate and its lipophilic analog BPH-703, bind in the allylic substrate (DMAPP, GPP, or FPP) site in the *Plasmodium vivax* FPPS/GGPPS homolog (No et al., 2012). This binding is similar to what is observed in bisphosphonate-mammalian FPPS complexes (Kavanagh et al., 2006; Rondeau et al., 2006; Yokoyama et al., 2015; Zhang et al., 2013). To determine whether MMV019313 and bisphosphonates share a common binding site, we compared the effect of the S228T variant on inhibition by MMV019313 and the bisphosphonate BPH-703. MMV019313 and BPH-703 both inhibited wild-type *Pf*FPPS/GGPPS activity. However only MMV019313 inhibition was decreased by over 10-fold in the S228T variant (Figure 4A-D). This difference in their *in vitro* enzyme inhibition was consistent with the effect on parasite growth inhibition, in which moderate overexpression of the S228T variant conferred resistance to MMV019313 but not BPH-703 (Figure S5). In addition, residue S228 is adjacent to the IPP binding site in the bisphosphonate-*Pv*FPPS/GGPPS structures and distal from the bound bisphosphonate. These results demonstrate that MMV019313 has a novel binding mode, distinct from the known allylic substrate site occupied by bisphosphonates, one which may confer its specificity for *Plasmodium* FPPS/GGPPS over human homologs.

Because our biochemical results indicated that MMV019313 binds a new site which is unique to *Pf*FPPS/GGPPS, we performed molecular docking calculations to evaluate the likelihood of MMV019313 binding at known small molecule binding sites in *Pf*FPPS/GGPPS. Three small molecule binding sites have been identified in FPPS and GPPS enzymes: 1) the allylic substrate binding site, which is also occupied by bisphosphonates, 2) the IPP binding site, and 3) an allosteric site identified in *Hs*FPPS (Jahnke et al., 2010). Because both the allylic substrate and IPP sites have been structurally characterized in *Pv*FPPS/GGPPS, we used the rigid body docking program Glide to generate poses with MMV019313 bound to these sites and estimate the binding free energy of each pose (Friesner et al., 2004; 2006; Halgren et al., 2004). The binding affinity estimated for the MMV019313 poses were >10^8^ weaker than bisphosphonate at the allylic site and >10^5^ weaker than IPP at its site (Table S4). Therefore, the modeling suggests that MMV019313 is unlikely to bind either the allylic substrate or IPP site. We also attempted to dock MMV019313 into other potential binding pockets in the *Pv*FPPS/GGPPS apo structure but with inconclusive results.

## Discussion

Farnesyl and geranylgeranyl diphosphate synthase (FPPS and GGPPS) are key branch point enzymes in isoprenoid biosynthesis. Human cells contain separate FPPS and GGPPS enzymes. An important class of clinical drugs, nitrogen-containing bisphosphonates, inhibits human FPPS in osteoclasts and block their function and proliferation (Kavanagh et al., 2006). Because osteoclasts are responsible for bone resorption, bisphosphonates are highly effective for treatment of osteoporosis and other bone remodeling diseases. Bisphosphonates are chemically stable analogs of inorganic pyrophosphate containing a P-C-P bond in place of the phosphodiester, which accounts for both its inhibition of FPPS (acting as an analog of the allylic diphosphate substrate) and its high selectivity for osteoclasts (depositing in bone mineral which is composed of calcium and phosphate). Unfortunately the charge state of bisphosphonates is a major liability in other therapeutic applications, as they are poorly bioavailable, rapidly cleared by the kidney, and do not achieve therapeutic levels in serum for treatment of non-bone diseases (Cremers et al., 2005; Jordão et al., 2011; Singh et al., 2010; Van Voorhis et al., 2016).

*Pf*FPPS/GGPPS, the molecular target of MMV019313 as demonstrated in this study, closely resembles mammalian FPPS enzymes in sequence, structure, and inhibition by bisphosphonates (Jordão et al., 2013; No et al., 2012). Like human FPPS, it is a central node in cellular isoprenoid biosynthesis vulnerable to drug inhibition (Artz et al., 2011; Kavanagh et al., 2006; Luckman et al., 1998). In *Plasmodium*, FPP and GGPP are required for the biosynthesis of prenylated proteins, the prenyl modification of ubiquinone, and other isoprenoid products, such that inhibition of *Pf*FPPS/GGPPS disrupts multiple cellular pathways (Chakrabarti et al., 2002; de Macedo et al., 2002; Gabriel et al., 2015; Tonhosolo et al., 2005; 2009). Previously lipophilic bisphosphonates modified with an alkyl chain to increase their cell permeability were shown to inhibit *Pv*FPPS/GGPPS homolog and clear both blood- and liver-stage *Plasmodium* parasites in mice infection models (No et al., 2012; Singh et al., 2010). Importantly, these results validated *Plasmodium* FPPS/GGPPS as an antiparasitic drug target for both acute malaria treatment and malaria chemoprophylaxis.

Our identification of MMV019313 further addresses two key hurdles in the development of *Pf*FPPS/GGPPS inhibitors as antimalarial drugs. First, MMV019313 represents the first non-bisphosphonate class of *Plasmodium* FPPS/GGPPS inhibitors with superior physicochemical properties. Many efforts have been made to develop modified bisphosphonates or non-bisphosphonate compounds as FPPS and/or GGPPS inhibitors for treatment of soft-tissue cancers and infectious diseases (Chen et al., 2013; Jahnke et al., 2010; Liu et al., 2014; Marzinzik et al., 2015; Zhang et al., 2009). Unlike bisphosphonates, MMV019313 has drug-like physicochemical properties satisfying the Rule of 5 and does not need to mimic a charged diphosphate substrate to achieve FPPS/GGPPS inhibition (Van Voorhis et al., 2016). As a compound in the Malaria Box library, it has already been tested in a panel of bioactivity and pharmacokinetic assays with encouraging results (Van Voorhis et al., 2016). Furthermore the results of >300 assays characterizing Malaria Box compounds as part of an innovative “open source” drug discovery effort by many groups will be a rich source of information (Allman et al., 2016; Paul et al., 2016; Ullah et al., 2017; Van Voorhis et al., 2016).

Second, MMV019313 has high selectivity for *Pf*FPPS/GGPPS over human FPPS and GGPPS, minimizing the potential for mechanism-based (e.g. “on-target”) toxicity. In fact, we could not detect any inhibition of human FPPS or GGPPS indicating selectivity at the enzymatic level of at least 100-fold. Consistent with this lack of enzymatic inhibition, MMV019313 showed no cytotoxity against a panel of 60 human cancer cell lines at 10 µM (Van Voorhis et al., 2016). In contrast, the most selective bisphosphonate BPH-703 identified by No *et al* showed 2.6-3.6-fold selectivity in our enzymatic assays (corresponding to a reported therapeutic index of 193 in growth inhibition assays) (No et al., 2012). Our results suggest that MMV019313 binds a novel site in *Pf*FPPS/GGPPS that is either absent from or substantially different in the human homologs, which may explain its high selectivity. Altogether MMV019313 offers a distinct and promising starting point for development of antimalarial FPPS/GGPPS inhibitors, which circumvents the inherent liabilities of the bisphosphonate pharmacophore and greatly improves on their selectivity.

Our immediate priority is to obtain structures of the inhibitor-enzyme complex to aid in structure-based design of MMV019313 derivatives with higher potency, as well as discovery of additional chemical scaffolds that occupy this new binding site. A potential challenge is that the FPPS/GGPPS structure is dynamic during catalysis. For example, the human FPPS is known to undergo at least two critical conformational changes (Kavanagh et al., 2006): The first is from an “open” apo form to a “closed” conformation upon binding of the allylic substrate, which is associated with drastically increased affinity for the allylic substrate (and bisphosphonates that mimic this substrate) as well as orders the IPP binding pocket. The second structural change occurs upon IPP binding and sequesters the active site from bulk solvent. Several attempts to obtain co-crystal structures of MMV019313 with *Pv*FPPS/GGPPS under conditions in which bisphosphonate-*Pv*FPPS/GGPPS crystals were obtained (presumed to correspond to the “closed” conformation) have been unsuccessful (personal communication, Dr. Raymond Hui). Additional crystallization conditions may require testing in the presence and absence of different allylic substrates and IPP, as their binding could affect the binding of MMV019313. Moreover the conformational dynamics of FPPS and GGPPS enzymes preclude straightforward interpretation of the mode-of-inhibition from conventional kinetic assays. Bisphosphonates that bind at the allylic substrate site show slow, tight binding with characteristics of irreversible inhibitors rather than classic competitive inhibitors (Dunford et al., 2008; Taylor, 2004). Indeed our preliminary assays indicate that MMV019313 also shows slow binding likely dependent on conformational changes.

Since the S228T variant distinguishes between MMV019313 and bisphosphonate binding to *Pf*FPPS/GGPPS, obtaining its structure to compare with wildtype FPPS/GGPPS will also offer clues to this new mode-of-inhibition. The resistance caused by the S228T variant could be explained by a direct contact between Ser228 and MMV019313 in a new small molecule binding pocket. But because the change from Ser to Thr is quite conservative, the addition of a methyl group, it seems more likely that Ser228 is involved in conformational dynamics important for catalysis. Structural analysis of this variant enzyme may reveal altered conformational states underlying the resistance to MMV019313.

Optimization of the bioactivity and pharmacokinetic properties of MMV019313 derivatives will be priorities for their development as antimalarials. The luciferase-based enzymatic assays performed in our study are robust for detection of effective inhibitors and can be adapted to high-throughput screening of small molecule libraries (in contrast we found a commercially-available colorimetric assay was not sufficiently sensitive) (Crowther et al., 2011). Inhibitor-enzyme structures will greatly accelerate optimization for potency against the *Plasmodium* FPPS/GGPPS, while simultaneously minimizing binding to the human homologs. *P. falciparum* strains overexpressing wildtype or the S228T variant, generated during this study, can also be re-tooled as secondary cellular assays for on-target specificity. Increased potency against blood-stage parasites will likely improve its activity against liver stage which currently is not detectable at 5 µM (Van Voorhis et al., 2016). Inhibition of isoprenoid biosynthesis by fosmidomycin, bisphosphonates, or MMV019313 has not been shown to block gametocyte development (Lell et al., 2003; Van Voorhis et al., 2016). The drug properties of MMV019313 derivatives will be optimized in standard absorption-distribution-metabolism-excretion-toxicity (ADME-T) assays. For example, we found that while MMV019313 is stable to human liver microsomal enzymes (*t*_½_> 159 min), its *t*_½_ in mouse liver microsomes was 4 min. This metabolic instability may account for a <1µM peak serum concentration following oral administration in mice and will need to be addressed before derivatives can be tested in mouse models of *Plasmodium* infection (Van Voorhis et al., 2016).

Finally, our study demonstrates two methodological improvements. First, we took advantage of a quantitative screen for IPP chemical rescue using a range of inhibitor concentrations, rather than a single high dose. The increased sensitivity allowed us to identify MMV019313, though it was missed in prior screens. It also permits the screen to be performed with geranylgeraniol, which is significantly less costly than IPP, since so far all identified inhibitors that rescue with IPP also show at least a partial rescue of parasite growth inhibition with geranylgeraniol (Yeh and Derisi, 2011; Zhang et al., 2011). Second, we successfully identified a drug-resistant mutation in chemically mutagenized *Plasmodium* parasites to facilitate mechanism-of-action elucidation. Though the rate and type of mutations caused by EMS was not characterized in our experiments, in the related apicomplexan parasite, *Toxoplasma gondii* EMS preferentially mutated A/T residues with similar rates of transitions and transversions (Farrell et al., 2014). Notably our study identified a T-to-A transversion in *Pf*FPPS/GGPPS in MMV019313-resistant parasites which were only selected after chemical mutagenesis. Characterization of the rate and type of mutations in *Plasmodium* parasites will facilitate selection and dosing of specific chemical mutagens to optimize phenotype selection and mutation identification by sequencing. Though limited, our experience suggests that the use of chemical mutagens will increase successful selection for drug resistance in *Plasmodium* parasites, which is the most common method to identify drug targets from phenotypic drug screens (Gamo et al., 2010; Guiguemde et al., 2010; Rottmann et al., 2010). The low frequency of *in vitro* MMV019313 resistance also indicates that clinical resistance will also be less frequent to MMV019313 and its derivatives.

### Significance

There is an urgent need for antimalarials with novel mechanisms-of-action to circumvent resistance to frontline drugs. Isoprenoid biosynthetic pathways are essential for *Plasmodium* (malaria) parasites and contain multiple validated drug targets. Using a chemical genetic approach from a phenotypic growth rescue screen to target identification, we identified a new antimalarial compound that inhibits a key branchpoint enzyme in isoprenoid biosynthesis. Our compound has significant advantages over bisphosphonate inhibitors that inhibit the same drug target with improved drug properties and specificity for the *Plasmodium* enzyme over human homologs. Our findings uncover a new “druggable” site in this important antimalarial drug target and provide a chemical starting point for development of *Plasmodium*-specific FPPS/GGPPS inhibitors.

## Materials and Methods

### P. *falciparum in vitro* cultures

*Plasmodium falciparum* W2 (MRA-157) and Dd2^attB^ (MRA-843) were obtained from MR4. Parasites were grown in human erythrocytes (2% hematocrit, obtained from the Stanford Blood Center) in RPMI 1640 media supplemented with 0.25% Albumax II (GIBCO Life Technologies), 2 g/L sodium bicarbonate, 0.1 mM hypoxanthine, 25 mM HEPES (pH 7.4), and 50 µg/L gentamycin, at 37°C, 5% O_2_, and 5% CO_2_. For passage of drug-treated, IPP-rescued parasites, the media was supplemented with 5 µM drug and 200 µM IPP (Isoprenoids LC or NuChem). For comparison of growth between different treatment conditions, cultures were carried simultaneously and handled identically with respect to media changes and addition of blood cells

### Chemical handling

Malaria Box compounds were received as 10 mM DMSO stocks in 96-well plates and diluted three-fold manually in DMSO. Fosmidomycin was included in control wells. Two-fold serial dilutions of the 96-well plates were performed on Velocity11. Compound stocks stored in DMSO were diluted for growth assays. MMV019313 was purchased from ChemDiv Biotech.

### qHTS screen for IPP chemical rescue

Growth assays were performed in 384-well clear bottom assay plates (E and K scientific, Santa Clara, CA). Drug (100-400 nL) was added directly to each well using PinTool (V&P Scientific) on a Sciclone ALH3000 (Caliper Sciences). Using the Titertek Multidrop 384, first 40 µL of growth media with and without 375 µM IPP was dispensed, followed by 10 µL ring-stage *P. falciparum* D10 parasites (parasitemia 0.8% in 10% hematocrit) into 384-well plates using the Titertek Multidrop 384. The final assay consisted of 50 µL ring-stage cultures at 0.8% parasitemia/2% hematocrit and drug concentrations from 0.01-26.7 µM ± 300 µM IPP. The plate was incubated at 37°C for 72 h. Parasites were lysed with 10 µL 5mM EDTA, 1.6% Triton-X, 20mM Tris-HCl and 0.16% Saponin containing 0.1% Sybr Green I (Invitrogen). The plates were then incubated at −80 °C for 20 min and thawed at room temperature overnight in the dark. Fluorescence was detected using Flexstation II-384. Compounds that showed IPP rescue of growth inhibition at 1 or more drug concentrations in the initial 384-well high-throughput screen were re-tested in a 96-well growth assay.

### Growth inhibition assays to determine EC_50_ values

*P. falciparum* cultures (125 µL) were grown in 96-well plates containing serial dilution of drugs in triplicate. Media was supplemented with 200 µM IPP as indicated. Growth was initiated with ring-stage parasites at 1% parasitemia and 0.5% hematocrit. Plates were incubated for 72h. Growth was terminated by fixation with 1% formaldehyde and parasitized cells were stained with 50 nM YOYO-1 (Invitrogen). Parasitemia was determined by flow cytometry. Data were analyzed by BD C6 Accuri C-Sampler software and fitted to a sigmoidal dose-inhibition function (Inhibition=Bottom + (Top-Bottom)/(1+10^((LogIC50-[drug])*HillSlope) by GraphPad Prism.

### *P. falciparum* mutagenesis and resistance selection

Late-stage parasites were purified using a SuperMACS II separator (Miltenyi Biotec) and incubated in complete medium with 8.3 – 2025 µM ethyl methanesulfonate (EMS, 6 concentrations total) for 2 hours. The concentrations of EMS used were selected by determining the EC_50_ in 72h parasite growth inhibition assays. The highest concentration used for mutagenesis was equal to the EC_50_ in order to maximize the selection pressure. The mutagen was then serially diluted 1:3 to test lower concentrations that might give lower mutation rates and therefore a lower frequency of passenger mutations. Mutagenized parasites were washed and separated into wells of 10 mL total volume (approximately 10% parasitemia, 2 % HCT). MMV019313 drug selection was applied to one well for each mutagenesis condition at 600 nM (approximately EC_75_), while the other well was left untreated in order to serve as a control for whole genome sequencing. Parasites were fed daily for the first week and every 3 days thereafter. Each culture was split in half every 6 days in order to introduce fresh RBC. Drug pressure was maintained for 32 days, with no observable parasite growth. After 32 days of selection, half of the culture from each EMS condition was removed from drug pressure. In these cultures, parasites which showed resistance to MMV019313 by a standard drug assay were observable after 7 days at all EMS conditions used. The parasites treated with the two lowest concentrations of EMS were selected for whole genome sequencing.

### Whole Genome Sequencing

*Plasmodium falciparum* strains were sequenced by Illumina sequencing as described previously (Straimer et al., 2012). Briefly, NEBNext DNA library reagents (NEB) and NEXTflex DNA barcode adapters (Bioo Scientific) were used to prepare PCR-free libraries (Kozarewa et al., 2009). Eight whole genome gDNA libraries were multiplexed and spiked with 8% PhiX control. Single-end sequencing was performed across two lanes on an Illumina HiSeq 2500 system. Data was analyzed using tools available in the Galaxy platform (Blankenberg et al., 2010; Giardine et al., 2005; Goecks et al., 2010). Sequencing reads were mapped against the *P. falciparum* 3D7v.10.0 reference genome using the Burrows-Wheeler Alignment tool (Li and Durbin, 2009). Sequencing data was visualized using Integrative Genomics Viewer (IGV) (Robinson et al., 2011; Thorvaldsdóttir et al., 2013). Variants were called using Freebayes (Garrison and Marth) and filtered for Quality >100 and Read Depth >30 using GATK tools (Van der Auwera et al., 2013). SnpEff was used to annotate the list of variants based on the *P. falciparum* 3D7v9.1 reference genome (Cingolani et al., 2012). Sequencing data have been deposited to the NCBI SRA (www.ncbi.nlm.nih.gov/sra; SRP106479).

### *P. falciparum* Transfections

An *E. coli* codon optimized version of *Pf*FPPS/GGPPS (PF3D7_1128400) was designed and synthesized by GeneWiz. Quick change mutagenesis was used to mutate GGPPS serine 228 to a threonine in the pUC vector provided by GeneWiz. These constructs were then moved into the pLN transfection plasmid designed for Bxb1 mycobacteriophage integrase system (Adjalley et al., 2010). The In-fusion cloning kit (Clontech) was used for all cloning. Restriction enzymes *Av*rII and *Bs*iWI were used to linearize the pLN vector. GGPPS was designed to have a *C*-terminal GFP tag. All transgenes were driven with either the ribosomal L2 protein (RL2) promoter (PF3D7_1132700) or the calmodulin (CaM) promoter (PF3D7_1434200) as denoted.

Transfections were carried out as previously described (Spalding et al., 2010). Briefly, 400 μL fresh red blood cells were preloaded with 100 μg of both pINT, which carries the bacteriophage integrase, and pRL2, which carries the gene of interest and the blasticidin resistance cassette, using a BioRad Gene-Pulser Xcell Electroporator. Electroporation conditions were infinite resistance, 310 V, and 950 μF using a 2 mm cuvette. Preloaded red blood cells were combined with 2.5 mL ~20% schizont Dd2^attB^ parasites and allowed to recover for 2 days before selection pressure was applied. Transfected parasites were selected with 2.5 μg/mL blasticidin are were detectable by thin smear within 15 days. Integration was confirmed by PCR and identity of the transgene was confirmed by sanger sequencing.

### Immunoblots

Parasites expressing either ACP_L_-GFP or one of the FPPS/GGPPS constructs generated in this study were isolated by saponin lysis and resuspended in 1xNuPage LDS sample buffer (Invitrogen). Whole cell lysate was separated by SDS-PAGE using 4-12% Bis-Tris gels (Initrogen) and transferred to nitrocellulose using a Trans Turbo-blot (BioRad). Membranes were blocked with 3% BSA, probed with 1:5,000 monoclonal anti-GFP JL-8 (BioRad) overnight, washed, then probed with 1:10,000 IRDye 680LT goat-anti-mouse. The Western was imaged using a Oddysey Imager (LiCor Biosciences).

### Live microscopy

Infected red blood cells were treated with Hoescht to stain the nucleus. Single z-stack images were collected on an epifluorescence Nikon eclipse.

### Quantative PCR

Quantitative PCR was performed as previously published (Yeh and Derisi, 2011). Briefly, parasites from 200 µL of culture were isolated by saponin lysis followed by PBS wash to remove extracellular DNA. DNA was purified using DNeasy Blood and Tissue kit (Qiagen). Primers were designed to target genes found on the apicoplast or nuclear genome: *tufA* (apicoplast) 5′-GATATTGATTCAGCTCCAGAAGAAA-3′ / 5′-ATATCCATTTGTGTGGCTCCTATAA-3′ and *CHT1* (nuclear) 5′-TGTTTCCTTCAACCCCTTTT-3′ / 5′-TGTTTCCTTCAACCCCTTTT-3′. Reactions contained template DNA, 0.15 µM of each primer, and 0.75× LightCycler 480 SYBR Green I Master mix (Roche). PCR reactions were performed at 56°C primer annealing and 65°C template extension for 35 cycles on a Lightcycler 6500 (Roche). For each time point, the apicoplast:nuclear genome ratio of the fosmidomycin-treated positive control, chloramphenicol-treated negative control, or MMV019313-treated experiment were calculated relative to that of an untreated control collected at the same time.

### Recombinant protein purification

Full length constructs of *Pf*FPPS/GGPPS, *Hs*FPPS, and *Hs*GGPPS were cloned into pET28a with an n-terminal hexahistidine tag. The *P. falciparum* FPPS/GGPPS was codon optimized for expression in *E. coli* (GeneWiz). *Pf*FPPS/GGPPS was mutagenized (S228T) using quick change mutagenesis. When expressed in *E. coli, Pf*FPPS/GGPPS (wt and S228T) and *Hs*FPPS were toxic. Cultures were supplemented with 0.4% glucose and grown to OD_600_ of 0.8-1 and induced with 0.5 mM IPTG. *Hs*GGPPS was grown without supplementation to an OD_600_ of 0.8-1. All cultures were induced for 4 hours at 37 °C, then harvested. Cells were lysed in 20 mM HEPES pH 8.0, 150 mM NaCl, 2 mM MgCl_2_, and protease inhibitor cocktail using sonication. Cleared lysates were either mixed with Talon metal affinity resin (Clonetech) or purified over 5 ml HisTrap columns (GE Healthcare). His tagged protein was purified with a single step purification eluting with buffer with 300 mM imidazole. Proteins were dialyzed to remove imidazole and flash frozen.

### *In vitro* FPPS/GGPPS assays

FPPS/GGPPS activity was measured by monitoring pyrophosphate release using the Lonza PPiLight kit under kinetic conditions. Drug and either 20 µg/ml *Pf*FPPS/GGPPS, 40 µg/ml *Hs*GGPPS, or 100 µg/ml *Hs*FPPS protein were incubated for 30 min room temperature. The reaction was initiated by the addition of Lonza PPiLight kit reaction mix and substrates. Saturating substrate conditions were 100 µM GPP or FPP, and 200 µM IPP. Sub-saturation conditions were K_M_ conditions as shown in Figure S4. Luciferase activity was monitored over time using a BioTek plate reader. Reaction rates were calculated from the linear portion of the raw luminescence over time curves (R^2^ >0.9). Data were fitted to a sigmoidal dose-inhibition function (Inhibition=Bottom + (Top-Bottom)/(1+10^((LogIC50-[drug])*HillSlope) by GraphPad Prism software.

### MMV019313 ligand docking

Using a solved *Pv*FPPS/GGPPS structure (PDB: 3EZ3, zoledronate and IPP bound) as a receptor model protein preparation wizard was used to add back in missing hydrogens and side chains. All crystallographic waters were removed. Hydrogen bonds were calculated using Epik at pH 8 (±1). The protein structure was minimized using OPL3 force field. MMV019313 was docked using glide to receptor grids generated from the relevant crystallized small molecules.

## Author Contributions

JEG, ZH, LO, and EY conducted experiments. JEG, ZH, LO, ML, and EY were responsible for data analysis. JEG and EY designed experiments and wrote the paper. ML and EY supervised the project.

## Acknowledgements

We are grateful to Medicines for Malaria Ventures (MMV) for providing the Malaria Box compounds and making this valuable library freely available, as well as GlaxoSmithKline for their screening efforts that first identified MMV019313 (TCMDC-123889). We would like to thank Dr. Susmitha Suresh for performing drug screens and Dr. Felice Kelly for advice on how to chemically mutagenize our parasites for resistance selection. We are grateful to Dr. James Dunford (University of Oxford) for advice in developing the *in vitro* enzyme activity assays. We also would like to acknowledge Dr. Wei Zhu and Professor Eric Oldfield (University of Illinois, Urbana-Champaign) for *in vitro P. vivax* GGPPS activity assays and providing BPH-703. Funding support for this project was generously provided by the Stanford Consortium for Innovation, Design, Evaluation and Action (C-IDEA), NIH 1K08AI097239 (EY), NIH 1DP5OD012119 (EY), the Burroughs Wellcome Fund Career Award for Medical Scientists (EY), the Burroughs Wellcome Fund Investigators in Pathogenesis of Infectious Disease (PATH) Award (ML), and the Stanford School of Medicine Dean's Postdoctoral Fellowship (JEG). The authors declare no conflict of interest.

## References

Adjalley, S.H., Lee, M.C.S., and Fidock, D.A. (2010). A method for rapid genetic integration into *Plasmodium falciparum* utilizing mycobacteriophage Bxb1 integrase. Methods Mol. Biol. 634, 87–100.

Allman, E.L., Painter, H.J., Samra, J., Carrasquilla, M., and Llinás, M. (2016). Metabolomic Profiling of the Malaria Box Reveals Antimalarial Target Pathways. Antimicrob. Agents Chemother. 60, 6635–6649.

Amberg-Johnson, K., Ganesan, S.M., Lorenzi, H.A., Niles, J.C., and Yeh, E. (2017). A first-in-class inhibitor of parasite FtsH disrupts plastid biogenesis in human pathogens. bioRxiv 108910.

Artz, J.D., Wernimont, A.K., Dunford, J.E., Schapira, M., Dong, A., Zhao, Y., Lew, J., Russell, R.G.G., Ebetino, F.H., Oppermann, U., et al. (2011). Molecular characterization of a novel geranylgeranyl pyrophosphate synthase from *Plasmodium* parasites. J. Biol. Chem. 286, 3315–3322.

Blankenberg, D., Kuster, Von, G., Coraor, N., Ananda, G., Lazarus, R., Mangan, M., Nekrutenko, A., and Taylor, J. (2010). Galaxy: a web-based genome analysis tool for experimentalists. Curr Protoc Mol Biol Chapter 19, Unit19.10.1–Unit19.10.21.

Bowman, J.D., Merino, E.F., Brooks, C.F., Striepen, B., Carlier, P.R., and Cassera, M.B. (2014). Antiapicoplast and gametocytocidal screening to identify the mechanisms of action of compounds within the malaria box. Antimicrob. Agents Chemother. 58, 811–819.

Chakrabarti, D., Da Silva, T., Barger, J., Paquette, S., Patel, H., Patterson, S., and Allen, C.M. (2002). Protein farnesyltransferase and protein prenylation in *Plasmodium falciparum*. J. Biol. Chem.

Chen, S.-H., Lin, S.-W., Lin, S.-R., Liang, P.-H., and Yang, J.-M. (2013). Moiety-linkage map reveals selective nonbisphosphonate inhibitors of human geranylgeranyl diphosphate synthase. J Chem Inf Model 53, 2299–2311.

Cingolani, P., Patel, V.M., Coon, M., Nguyen, T., Land, S.J., Ruden, D.M., and Lu, X. (2012). Using *Drosophila melanogaster* as a Model for Genotoxic Chemical Mutational Studies with a New Program, SnpSift. Front Genet 3, 35.

Creek, D.J., Chua, H.H., Cobbold, S.A., Nijagal, B., MacRae, J.I., Dickerman, B.K., Gilson, P.R., Ralph, S.A., and McConville, M.J. (2016). Metabolomics-Based Screening of the Malaria Box Reveals both Novel and Established Mechanisms of Action. Antimicrob. Agents Chemother. 60, 6650–6663.

Cremers, S.C.L.M., Pillai, G., and Papapoulos, S.E. (2005). Pharmacokinetics/pharmacodynamics of bisphosphonates: use for optimisation of intermittent therapy for osteoporosis. Clin Pharmacokinet 44, 551–570.

Crowther, G.J., Napuli, A.J., Gilligan, J.H., Gagaring, K., Borboa, R., Francek, C., Chen, Z., Dagostino, E.F., Stockmyer, J.B., Wang, Y., et al. (2011). Identification of inhibitors for putative malaria drug targets among novel antimalarial compounds. Mol. Biochem. Parasitol. 175, 21–29.

de Macedo, C.S., Uhrig, M.L., Kimura, E.A., and Katzin, A.M. (2002). Characterization of the isoprenoid chain of coenzyme Q in *Plasmodium falciparum*. FEMS Microbiol. Lett. 207, 13–20.

DeRisi, J.L. (2014). UCSF DeRisi Lab MMV Box Apicoplast Screening (EMBL-EBI).

Desai, S.A., Alkhalil, A., Kang, M., Ashfaq, U., and Nguyen, M.-L. (2005). Plasmodial surface anion channel-independent phloridzin resistance in *Plasmodium falciparum*. J. Biol. Chem. 280, 16861–16867.

Dunford, J.E., Kwaasi, A.A., Rogers, M.J., Barnett, B.L., Ebetino, F.H., Russell, R.G.G., Oppermann, U., and Kavanagh, K.L. (2008). Structure-activity relationships among the nitrogen containing bisphosphonates in clinical use and other analogues: time-dependent inhibition of human farnesyl pyrophosphate synthase. J. Med. Chem. 51, 2187–2195.

Eastman, R.T., White, J., Hucke, O., Bauer, K., Yokoyama, K., Nallan, L., Chakrabarti, D., Verlinde, C.L.M.J., Gelb, M.H., Rathod, P.K., et al. (2005). Resistance to a protein farnesyltransferase inhibitor in *Plasmodium falciparum*. J. Biol. Chem. 280, 13554–13559.

Eastman, R.T., White, J., Hucke, O., Yokoyama, K., Verlinde, C.L.M.J., Hast, M.A., Beese, L.S., Gelb, M.H., Rathod, P.K., and Van Voorhis, W.C. (2007). Resistance mutations at the lipid substrate binding site of *Plasmodium falciparum* protein farnesyltransferase. Mol. Biochem. Parasitol. 152, 66–71.

Farrell, A., Coleman, B.I., Benenati, B., Brown, K.M., Blader, I.J., Marth, G.T., and Gubbels, M.-J. (2014). Whole genome profiling of spontaneous and chemically induced mutations in *Toxoplasma gondii*. BMC Genomics 15, 354.

Fernandes, J.F., Lell, B., Agnandji, S.T., Obiang, R.M., Bassat, Q., Kremsner, P.G., Mordmüller, B., and Grobusch, M.P. (2015). Fosmidomycin as an antimalarial drug: a meta-analysis of clinical trials. Future Microbiol 10, 1375–1390.

Friesner, R.A., Banks, J.L., Murphy, R.B., Halgren, T.A., Klicic, J.J., Mainz, D.T., Repasky, M.P., Knoll, E.H., Shelley, M., Perry, J.K., et al. (2004). Glide: a new approach for rapid, accurate docking and scoring. *1. Method and assessment of docking accuracy*. J. Med. Chem. 47, 1739–1749.

Friesner, R.A., Murphy, R.B., Repasky, M.P., Frye, L.L., Greenwood, J.R., Halgren, T.A., Sanschagrin, P.C., and Mainz, D.T. (2006). Extra precision glide: docking and scoring incorporating a model of hydrophobic enclosure for protein-ligand complexes. J. Med. Chem. 49, 6177–6196.

Gabriel, H.B., Silva, M.F., Kimura, E.A., Wunderlich, G., Katzin, A.M., and Azevedo, M.F. (2015). Squalestatin is an inhibitor of carotenoid biosynthesis in *Plasmodium falciparum*. Antimicrob. Agents Chemother. 59, 3180–3188.

Gamo, F.J., Sanz, L.M., Vidal, J., de Cozar, C., Alvarez, E., Lavandera, J.-L., Vanderwall, D.E., Green, D.V.S., Kumar, V., Hasan, S., et al. (2010). Thousands of chemical starting points for antimalarial lead identification. Nature 465, 305–310.

Garrison, E., and Marth, G. Haplotype-based variant detection from short-read sequencing. Httpsarxiv.orgabs.V 2012.

Ghosh, S., Chan, J.M.W., Lea, C.R., Meints, G.A., Lewis, J.C., Tovian, Z.S., Flessner, R.M., Loftus, T.C., Bruchhaus, I., Kendrick, H., et al. (2004). Effects of bisphosphonates on the growth of *Entamoeba histolytica* and *Plasmodium* species in vitro and in vivo. J. Med. Chem. 47, 175–187.

Giardine, B., Riemer, C., Hardison, R.C., Burhans, R., Elnitski, L., Shah, P., Zhang, Y., Blankenberg, D., Albert, I., Taylor, J., et al. (2005). Galaxy: a platform for interactive large-scale genome analysis. Genome Res. 15, 1451–1455.

Goecks, J., Nekrutenko, A., Taylor, J., Galaxy Team (2010). Galaxy: a comprehensive approach for supporting accessible, reproducible, and transparent computational research in the life sciences. Genome Biol. 11, R86.

Guiguemde, W.A., Shelat, A.A., Bouck, D., Duffy, S., Crowther, G.J., Davis, P.H., Smithson, D.C., Connelly, M., Clark, J., Zhu, F., et al. (2010). Chemical genetics of *Plasmodium falciparum*. Nature 465, 311–315.

Halgren, T.A., Murphy, R.B., Friesner, R.A., Beard, H.S., Frye, L.L., Pollard, W.T., and Banks, J.L. (2004). Glide: a new approach for rapid, accurate docking and scoring. *2. Enrichment factors in database screening*. J. Med. Chem. 47, 1750–1759.

Inselburg, J. (1984). Induction and selection of drug resistant mutants of *Plasmodium falciparum*. Mol. Biochem. Parasitol. 10, 89–98.

Inselburg, J. (1985). Induction and isolation of artemisinine-resistant mutants of *Plasmodium falciparum*. Am. J. Trop. Med. Hyg. 34, 417–418.

Jahnke, W., Rondeau, J.-M., Cotesta, S., Marzinzik, A., Pellé, X., Geiser, M., Strauss, A., Götte, M., Bitsch, F., Hemmig, R., et al. (2010). Allosteric non-bisphosphonate FPPS inhibitors identified by fragment-based discovery. Nat. Chem. Biol. 6, 660–666.

Jomaa, H., Wiesner, J., Sanderbrand, S., Altincicek, B., Weidemeyer, C., Hintz, M., Türbachova, I., Eberl, M., Zeidler, J., Lichtenthaler, H.K., et al. (1999). Inhibitors of the nonmevalonate pathway of isoprenoid biosynthesis as antimalarial drugs. Science 285, 1573–1576.

Jordão, F.M., Gabriel, H.B., Alves, J.M.P., Angeli, C.B., Bifano, T.D., Breda, A., de Azevedo, M.F., Basso, L.A., Wunderlich, G., Kimura, E.A., et al. (2013). Cloning and characterization of bifunctional enzyme farnesyl diphosphate/geranylgeranyl diphosphate synthase from *Plasmodium falciparum*. Malar. J. 12, 184.

Jordão, F.M., Saito, A.Y., Miguel, D.C., de Jesus Peres, V., Kimura, E.A., and Katzin, A.M. (2011). *In vitro* and *in vivo* antiplasmodial activities of risedronate and its interference with protein prenylation in *Plasmodium falciparum*. Antimicrob. Agents Chemother. 55, 2026–2031.

Kavanagh, K.L., Guo, K., Dunford, J.E., Wu, X., Knapp, S., Ebetino, F.H., Rogers, M.J., Russell, R.G.G., and Oppermann, U. (2006). The molecular mechanism of nitrogen-containing bisphosphonates as antiosteoporosis drugs. Proc. Natl. Acad. Sci. U.S.a. 103, 7829–7834.

Kozarewa, I., Ning, Z., Quail, M.A., Sanders, M.J., Berriman, M., and Turner, D.J. (2009). Amplification-free Illumina sequencing-library preparation facilitates improved mapping and assembly of (G+C)-biased genomes. Nat. Methods 6, 291–295.

Lell, B., Ruangweerayut, R., Wiesner, J., Missinou, M.A., Schindler, A., Baranek, T., Hintz, M., Hutchinson, D., Jomaa, H., and Kremsner, P.G. (2003). Fosmidomycin, a novel chemotherapeutic agent for malaria. Antimicrob. Agents Chemother. 47, 735–738.

Li, H., and Durbin, R. (2009). Fast and accurate short read alignment with Burrows-Wheeler transform. Bioinformatics 25, 1754–1760.

Liu, J., Liu, W., Ge, H., Gao, J., He, Q., Su, L., Xu, J., Gu, L.-Q., Huang, Z.-S., and Li, D. (2014). Syntheses and characterization of non-bisphosphonate quinoline derivatives as new FPPS inhibitors. Biochim. Biophys. Acta 1840, 1051–1062.

Luckman, S.P., Hughes, D.E., Coxon, F.P., Graham, R., Russell, G., and Rogers, M.J. (1998). Nitrogen-containing bisphosphonates inhibit the mevalonate pathway and prevent post-translational prenylation of GTP-binding proteins, including Ras. J. Bone Miner. Res. 13, 581–589.

Marzinzik, A.L., Amstutz, R., Bold, G., Bourgier, E., Cotesta, S., Glickman, J.F., Götte, M., Henry, C., Lehmann, S., Hartwieg, J.C.D., et al. (2015). Discovery of Novel Allosteric Non-Bisphosphonate Inhibitors of Farnesyl Pyrophosphate Synthase by Integrated Lead Finding. ChemMedChem 10, 1884–1891.

Michael B Martin, Joshua S Grimley, Jared C Lewis, Huel T Heath, III, Brian N Bailey, Howard Kendrick, Vanessa Yardley, Aura Caldera, Renee Lira, Julio A Urbina, et al. (2001). Bisphosphonates Inhibit the Growth of Trypanosoma brucei, Trypanosoma cruzi, Leishmania donovani, Toxoplasma gondii, and Plasmodium falciparum: A Potential Route to Chemotherapy. J. Med. Chem. 44, 909–916.

Nallan, L., Bauer, K.D., Bendale, P., Rivas, K., Yokoyama, K., Hornéy, C.P., Pendyala, P.R., Floyd, D., Lombardo, L.J., Williams, D.K., et al. (2005). Protein farnesyltransferase inhibitors exhibit potent antimalarial activity. J. Med. Chem. 48, 3704–3713.

Nkrumah, L.J., Muhle, R.A., Moura, P.A., Ghosh, P., Hatfull, G.F., Jacobs, W.R., and Fidock, D.A. (2006). Efficient site-specific integration in *Plasmodium falciparum* chromosomes mediated by mycobacteriophage Bxb1 integrase. Nat. Methods 3, 615–621.

No, J.H., de Macedo Dossin, F., Zhang, Y., Liu, Y.-L., Zhu, W., Feng, X., Yoo, J.A., Lee, E., Wang, K., Hui, R., et al. (2012). Lipophilic analogs of zoledronate and risedronate inhibit *Plasmodium* geranylgeranyl diphosphate synthase (GGPPS) and exhibit potent antimalarial activity. Proc. Natl. Acad. Sci. U.S.a. 109, 4058–4063.

Otto, T.D., Wilinski, D., Assefa, S., Keane, T.M., Sarry, L.R., Böhme, U., Lemieux, J., Barrell, B., Pain, A., Berriman, M., et al. (2010). New insights into the blood-stage transcriptome of *Plasmodium falciparum* using RNA-Seq. Mol. Microbiol. 76, 12–24.

Paul, A.S., Moreira, C.K., Elsworth, B., Allred, D.R., and Duraisingh, M.T. (2016). Extensive Shared Chemosensitivity between Malaria and Babesiosis Blood-Stage Parasites. Antimicrob. Agents Chemother. 60, 5059–5063.

Robinson, J.T., Thorvaldsdóttir, H., Winckler, W., Guttman, M., Lander, E.S., Getz, G., and Mesirov, J.P. (2011). Integrative genomics viewer. Nat. Biotechnol. 29, 24–26.

Rondeau, J.-M., Bitsch, F., Bourgier, E., Geiser, M., Hemmig, R., Kroemer, M., Lehmann, S., Ramage, P., Rieffel, S., Strauss, A., et al. (2006). Structural Basis for the Exceptional in vivo Efficacy of Bisphosphonate Drugs. ChemMedChem 1, 267–273.

Rottmann, M., McNamara, C., Yeung, B.K.S., Lee, M.C.S., Zou, B., Russell, B., Seitz, P., Plouffe, D.M., Dharia, N.V., Tan, J., et al. (2010). Spiroindolones, a potent compound class for the treatment of malaria. Science 329, 1175–1180.

Singh, A.P., Zhang, Y., No, J.H., Docampo, R., Nussenzweig, V., and Oldfield, E. (2010). Lipophilic bisphosphonates are potent inhibitors of *Plasmodium* liver-stage growth. Antimicrob. Agents Chemother. 54, 2987–2993.

Sinigaglia, L., Varenna, M., and Casari, S. (2007). Pharmacokinetic profile of bisphosphonates in the treatment of metabolic bone disorders. Clin Cases Miner Bone Metab 4, 30–36.

Spalding, M.D., Allary, M., Gallagher, J.R., and Prigge, S.T. (2010). Validation of a modified method for Bxb1 mycobacteriophage integrase-mediated recombination in *Plasmodium falciparum* by localization of the H-protein of the glycine cleavage complex to the mitochondrion. Mol. Biochem. Parasitol. 172, 156–160.

Spangenberg, T., Burrows, J.N., Kowalczyk, P., McDonald, S., Wells, T.N.C., and Willis, P. (2013). The open access malaria box: a drug discovery catalyst for neglected diseases. PLoS ONE 8, e62906.

Straimer, J., Lee, M.C.S., Lee, A.H., Zeitler, B., Williams, A.E., Pearl, J.R., Zhang, L., Rebar, E.J., Gregory, P.D., Llinás, M., et al. (2012). Site-specific genome editing in *Plasmodium falciparum* using engineered zinc-finger nucleases. Nat. Methods 9, 993–998.

Taylor, K.B. (2004). Slow and Tight Inhibition. In Enzyme Kinetics and Mechanisms, (Dordrecht: Kluwer Academic Publishers), pp. 122–146.

Thorvaldsdóttir, H., Robinson, J.T., and Mesirov, J.P. (2013). Integrative Genomics Viewer (IGV): high-performance genomics data visualization and exploration. Brief. Bioinformatics 14, 178–192.

Tonhosolo, R., D’Alexandri, F.L., de Rosso, V.V., Gazarini, M.L., Matsumura, M.Y., Peres, V.J., Merino, E.F., Carlton, J.M., Wunderlich, G., Mercadante, A.Z., et al. (2009). Carotenoid biosynthesis in intraerythrocytic stages of *Plasmodium falciparum*. J. Biol. Chem. 284, 9974–9985.

Tonhosolo, R., D’Alexandri, F.L., Genta, F.A., Wunderlich, G., Gozzo, F.C., Eberlin, M.N., Peres, V.J., Kimura, E.A., and Katzin, A.M. (2005). Identification, molecular cloning and functional characterization of an octaprenyl pyrophosphate synthase in intra-erythrocytic stages of *Plasmodium falciparum*. Biochem. J. 392, 117–126.

Tsuchiya, T., Ishibashi, K., Terakawa, M., Nishiyama, M., Itoh, N., and Noguchi, H. (1982). Pharmacokinetics and metabolism of fosmidomycin, a new phosphonic acid, in rats and dogs. Eur J Drug Metab Pharmacokinet 7, 59–64.

Ullah, I., Sharma, R., Biagini, G.A., and Horrocks, P. (2017). A validated bioluminescence-based assay for the rapid determination of the initial rate of kill for discovery antimalarials. J. Antimicrob. Chemother. 72, 717–726.

Van der Auwera, G.A., Carneiro, M.O., Hartl, C., Poplin, R., Del Angel, G., Levy-Moonshine, A., Jordan, T., Shakir, K., Roazen, D., Thibault, J., et al. (2013). From FastQ data to high confidence variant calls: the Genome Analysis Toolkit best practices pipeline. Curr Protoc Bioinformatics 43, 11.10.1–.10.33.

Van Voorhis, W.C., Adams, J.H., Adelfio, R., Ahyong, V., Akabas, M.H., Alano, P., Alday, A., Alemán Resto, Y., Alsibaee, A., Alzualde, A., et al. (2016). Open Source Drug Discovery with the Malaria Box Compound Collection for Neglected Diseases and Beyond. PLoS Pathog. 12, e1005763.

Van Voorhis, W.C., Rivas, K.L., Bendale, P., Nallan, L., Hornéy, C., Barrett, L.K., Bauer, K.D., Smart, B.P., Ankala, S., Hucke, O., et al. (2007). Efficacy, pharmacokinetics, and metabolism of tetrahydroquinoline inhibitors of *Plasmodium falciparum* protein farnesyltransferase. Antimicrob. Agents Chemother. 51, 3659–3671.

Wu, W., Herrera, Z., Ebert, D., Baska, K., Cho, S.H., Derisi, J.L., and Yeh, E. (2015). A chemical rescue screen identifies a Plasmodium falciparum apicoplast inhibitor targeting MEP isoprenoid precursor biosynthesis. Antimicrob. Agents Chemother. 59, 356–364.

Yeh, E., and Derisi, J.L. (2011). Chemical rescue of malaria parasites lacking an apicoplast defines organelle function in blood-stage *Plasmodium falciparum*. PLoS Biol. 9, e1001138.

Yokoyama, T., Mizuguchi, M., Ostermann, A., Kusaka, K., Niimura, N., Schrader, T.E., and Tanaka, I. (2015). Protonation State and Hydration of Bisphosphonate Bound to Farnesyl Pyrophosphate Synthase. J. Med. Chem. 58, 7549–7556.

Zhang, B., Watts, K.M., Hodge, D., Kemp, L.M., Hunstad, D.A., Hicks, L.M., and Odom, A.R. (2011). A second target of the antimalarial and antibacterial agent fosmidomycin revealed by cellular metabolic profiling. Biochemistry 50, 3570–3577.

Zhang, Y., Cao, R., Yin, F., Hudock, M.P., Guo, R.-T., Krysiak, K., Mukherjee, S., Gao, Y.-G., Robinson, H., Song, Y., et al. (2009). Lipophilic bisphosphonates as dual farnesyl/geranylgeranyl diphosphate synthase inhibitors: an X-ray and NMR investigation. J. Am. Chem. Soc. 131, 5153–5162.

Zhang, Y., Zhu, W., Liu, Y.-L., Wang, H., Wang, K., Li, K., No, J.H., Ayong, L., Gulati, A., Pang, R., et al. (2013). Chemo-Immunotherapeutic Anti-Malarials Targeting Isoprenoid Biosynthesis. ACS Med Chem Lett 4, 423–427.

